# The Interaction of LILRB2 with HLA-B is Associated with Psoriasis Susceptibility

**DOI:** 10.1101/741983

**Authors:** Rebecca L. Yanovsky, Haoyan Chen, Stephen Leslie, Mary Carrington, Wilson Liao

## Abstract

Genetic variation within the major histocompatibility complex (MHC) class I is a well-known risk factor for psoriasis. While the mechanisms behind this variation are still being fully elucidated, human leukocyte antigen (HLA) presentation of auto-antigens as well as the interaction of HLA-B with killer cell immunoglobulin-like receptors (KIRs) have been shown to contribute to psoriasis susceptibility. Here we demonstrate that the interaction of HLA class I molecules with leukocyte immunoglobulin-like receptors (LILR), a related group of immunomodulatory receptors primarily found on antigen presenting cells, also contributes to psoriasis susceptibility. We used previously characterized binding capacities of HLA-A, HLA-B, and HLA-C allotypes to two inhibitory LILRs, LILRB1 and LILRB2, to investigate the effect of LILRB1/2 binding in two large genome wide association study cohorts of psoriasis patients and controls (N = 10,069). We found that the strength of binding of LILRB2 to HLA-B was inversely associated with psoriasis risk (p = 2.34E-09, OR [95% CI], 0.41 [0.30−0.55]) independent of individual class I or II allelic effects. We thus propose that weak binding of inhibitory LILRB2 to HLA-B may play a role in patient susceptibility to psoriasis via increased activity of antigen presenting cells.

## INTRODUCTION

The strongest signal for genetic susceptibility to psoriasis resides within the major histocompatibility complex (MHC), particularly near MHC class I loci (Okada et al., 2014). However, the mechanistic basis for how these MHC associations confer psoriasis risk has yet to be fully elucidated. Fine-mapping studies in this region have identified multiple independent effects for various class I alleles, with specific amino acids in the peptide binding groove of HLA-B and HLA-C being important (Chen et al., 2012, Okada et al., 2014). This suggests that psoriasis risk may be mediated through HLA presentation of auto-antigens. However, MHC class I have other important immunoregulatory functions, including binding and regulation of natural killer (NK) cells through killer cell immunoglobulin-like receptors (KIRs) or regulation of antigen presenting cells (APCs) through leukocyte immunoglobulin-like receptors (LILR). We have previously shown that HLA-B can mediate psoriasis risk through interaction with KIRs (Ahn et al., 2016). Here, we demonstrate that HLA-B can also mediate psoriasis susceptibility through differential binding to LILRs.

LILRs are a cousin of KIRs, closely located on chromosome 19, but are primarily found on APCs such as dendritic cells and macrophages, as well as subsets of B-cells, T-cells, and NK cells (Jones et al., 2011). LILRs participate in regulation of APC function through engagement of either activating receptors (LILRA) or inhibitory receptors (LILRB). LILRA increase secretion of inflammatory cytokines and basophil degranulation by increasing monocyte activation (Hudson and Allen, 2016). Conversely, LILRB inhibit co-stimulatory proteins on APCs, may reduce antigen presentation on these cells, and facilitate an increased regulatory T cell response (Bashirova et al., 2014, Hudson and Allen, 2016).

LILR have been associated with several autoimmune and infectious diseases (Zhang et al., 2017). To further elucidate the basis of psoriasis susceptibility due to HLA alleles, we examined the distribution of binding affinities of LILRB1 and LILRB2 to HLA-A, -B, and -C in two large psoriasis genome-wide association study (GWAS) cohorts totaling 10,069 subjects.

## RESULTS

We examined the role of LILRB1 and LILRB2 in psoriasis by testing six variables: LILRB1 or LILRB2 binding to HLA-A, -B, and -C. In the Wellcome Trust Case-Control Consortium (WTCCC) discovery cohort, stepwise regression of these six variables along with inclusion of all individual imputed HLA class I and class II alleles, adjusted for gender and the first ten principal components of ancestry, revealed a number of independently associated variables (**Table 1**). As expected, the strongest signal was for HLA-C*06:02. However, we also identified an association between the binding strength of LILRB2 to HLA-B (LILRB2-B) with an OR [95% CI] of 0.44 [0.28−0.68], p = 2.33E-04. This OR being < 1 implies stronger LILRB2-B binding decreases the risk for psoriasis; alternatively stated, weaker binding increases risk for psoriasis. No other LILRB variable demonstrated significance at the level of p<0.01. In a second, independent cohort (Genetic Association Information Network (GAIN)), we validated that reduced binding of LILRB2 to HLA-B allotypes promotes psoriasis (p=3.43E-03, OR 0.50 [0.32−0.80]), (**Table 1**).

**Table 1.** Stepwise regression analysis of the association of LILRB binding and individual HLA alleles with psoriasis in the WTCCC and GAIN cohorts

| **WTCCC (Discovery Cohort)** |  |  | **GAIN (Replication Cohort)** |  |  |
| --- | --- | --- | --- | --- | --- |
| **2,178 cases, 5,175 controls** |  |  | **1,368 cases, 1,348 controls** |  |  |
| **Variable** | **p-value** | **OR (95% CI)** | **Variable** | **p-value** | **OR (95% CI)** |
| C0602 | 3.32E-26 | 4.20 (3.22-5.47) | C0602 | 5.27E-35 | 4.41 (3.49-5.58) |
| A0201 | 1.83E-13 | 1.70 (1.47-1.95) | DQB0604 | 2.65E-06 | 0.37 (0.25-0.56) |
| A0101 | 4.64E-08 | 1.62 (1.36-1.92) | B5001 | 2.24E-03 | 0.43 (0.25-0.74) |
| B3801 | 2.78E-02 | 1.82 (1.07-3.11) | B3801 | 3.37E-04 | 2.20 (1.43-3.39) |
| B5501 | 5.41E-05 | 1.93 (1.40-2.65) | A0201 | 1.32E-03 | 1.34 (1.12-1.61) |
| **LILRB2-B** | **2.33E-04** | **0.44 (0.28-0.68)** | B3901 | 3.17E-03 | 2.08 (1.28-3.37) |
| DQB0604 | 1.30E-02 | 0.64 (0.45-0.91) | DRB0801 | 6.81E-03 | 0.56 (0.37-0.85) |
| A1101 | 1.64E-02 | 1.31 (1.05-1.62) | B5501 | 3.47E-03 | 1.91 (1.24-2.95) |
| C1203 | 2.66E-02 | 1.54 (1.05-2.26) | B0801 | 2.10E-03 | 1.45 (1.14-1.83) |
| B1302 | 6.15E-04 | 1.74 (1.27-2.38) | **LILRB2-B** | **3.43E-03** | **0.50 (0.32-0.80)** |
| B5701 | 1.41E-02 | 1.49 (1.08-2.04) |  |  |  |
| B4001 | 2.69E-02 | 0.75 (0.58-0.97) |  |  |  |
| DQA0501 | 1.88E-03 | 0.77 (0.65-0.91) |  |  |  |
| B0801 | 7.66E-03 | 1.36 (1.08-1.70) |  |  |  |
| B4403 | 4.09E-02 | 0.77 (0.59-0.99) |  |  |  |

Abbreviations: GWAS, Genome Wide Association Studies; WTCCC, Wellcome Trust Case-Control Consortium.

Joint analysis of the WTCCC and GAIN cohorts indicated a significant effect of LILRB2-B binding level in both stepwise regression analysis (p=2.20E-09, OR 0.41 [0.30−0.55]) and multivariate analysis (p=2.34E-09, OR 0.41 [0.30−0.55]) **(Table 2)**. These results indicate that psoriasis is associated with weaker binding of HLA-B to LILRB2, independent of class I or II allelic effects.

**Table 2.** Stepwise regression and multivariate association analysis of LILRB binding and individual HLA alleles with psoriasis in the combined WTCCC and GAIN cohorts. All variables from the stepwise analysis were included in the multivariate analysis.

| Variable | Stepwise Univariate Analysis |  | Multivariate Analysis |  |
| --- | --- | --- | --- | --- |
|  | p-value | OR (95% CI) | p-value | OR (95% CI) |
| C0602 | 1.71E-123 | 5.29 (4.61-6.08) | 1.33E-123 | 5.30 (4.62-6.09) |
| B5001 | 1.71E-06 | 0.45 (0.33-0.62) | 1.83E-06 | 0.45 (0.33-0.63) |
| A0201 | 2.51E-15 | 1.57 (1.40-1.75) | 2.32E-15 | 1.57 (1.40-1.75) |
| A0101 | 3.68E-07 | 1.43 (1.24-1.63) | 4.45E-07 | 1.42 (1.24-1.63) |
| DQB0604 | 5.53E-06 | 0.53 (0.40-0.70) | 4.94E-06 | 0.53 (0.40-0.69) |
| B5501 | 2.39E-06 | 1.84 (1.43-2.37) | 2.57E-06 | 1.84 (1.43-2.37) |
| **LILRB2-B** | **2.20E-09** | **0.41 (0.30-0.55)** | **2.34E-09** | **0.41 (0.30-0.55)** |
| B4001 | 3.58E-03 | 0.75 (0.62-0.91) | 3.53E-03 | 0.75 (0.62-0.91) |
| B4403 | 1.81E-03 | 0.73 (0.60-0.89) | 1.83E-03 | 0.73 (0.60-0.89) |
| C1203 | 4.29E-07 | 1.72 (1.40-2.13) | 3.59E-07 | 1.73 (1.40-2.14) |
| B0801 | 5.40E-05 | 1.75 (1.34-2.30) | 6.54E-05 | 1.74 (1.33-2.29) |
| DQA0501 | 1.35E-02 | 0.85 (0.75-0.97) | 1.44E-02 | 0.85 (0.75-0.97) |
| A1101 | 3.61E-02 | 1.20 (1.01-1.43) | 3.87E-02 | 1.20 (1.01-1.43) |
| C0701 | 3.09E-02 | 0.76 (0.60-0.98) | 3.42E-02 | 0.77 (0.60-0.98) |
| DRB0801 | 4.76E-02 | 0.75 (0.56-1.00) | 5.47E-02 | 0.75 (0.56-1.01) |

Abbreviations: GWAS, Genome Wide Association Studies; WTCCC, Wellcome Trust Case-Control Consortium.

To understand the cellular localization of LILRB2 in human skin, we examined LILRB2 gene expression in FACS-sorted keratinocytes, myeloid dendritic cells, and T lymphocytes from the skin of healthy human subjects (Ahn et al., 2017). We found robust expression of LILRB2 in cutaneous dendritic cells and negligible expression in keratinocytes and T cells (**Figure 1**). We also found that LILRB2 is significantly overexpressed (p < 5.0E-08, fold change=2.2) in psoriasis lesional skin (n=58) compared to healthy skin (n=64) (**Table 3**), consistent with the known increased number of dendritic cells in psoriatic skin. We have previously shown that the ability of monocyte-derived dendritic cells to induce proliferation of allogeneic CD4 T cells after exposure to a panel of recombinant class I molecules is inversely proportional to binding scores of the HLA allotype to LILRB2; whereby weak binding to LILRB2 results in high proliferation and strong binding to LILRB2 results in lower proliferation (Bashirova et al., 2014). CD4 T cell proliferation in this mixed leukocyte reaction was enhanced by the addition of LILRB2 siRNA (Bashirova et al., 2014). Thus, while dendritic cells expressing LILRB2 are commonly observed in skin, their ability to enhance T cell proliferation may be diminished in the presence of HLA allotypes that bind strongly to inhibitory LILRB2.

**Table 3.** *LILRB2* mRNA is significantly overexpressed in psoriasis lesional skin compared to healthy skin.

| Gene | Probe ID | Adjusted P Value | Fold Change<br>(Psoriasis lesional (n=58) vs. healthy skin (n=64)) |
| --- | --- | --- | --- |
| *LILRB2* | 210146_x_at | 3.84E-10 | 2.39 |
| *LILRB2* | 207697_x_at | 1.04E-09 | 2.20 |
| Source: https://www.ncbi.nlm.nih.gov/geo/query/acc.cgi?acc=GSE13355 |  |  |  |

**Figure 1.**
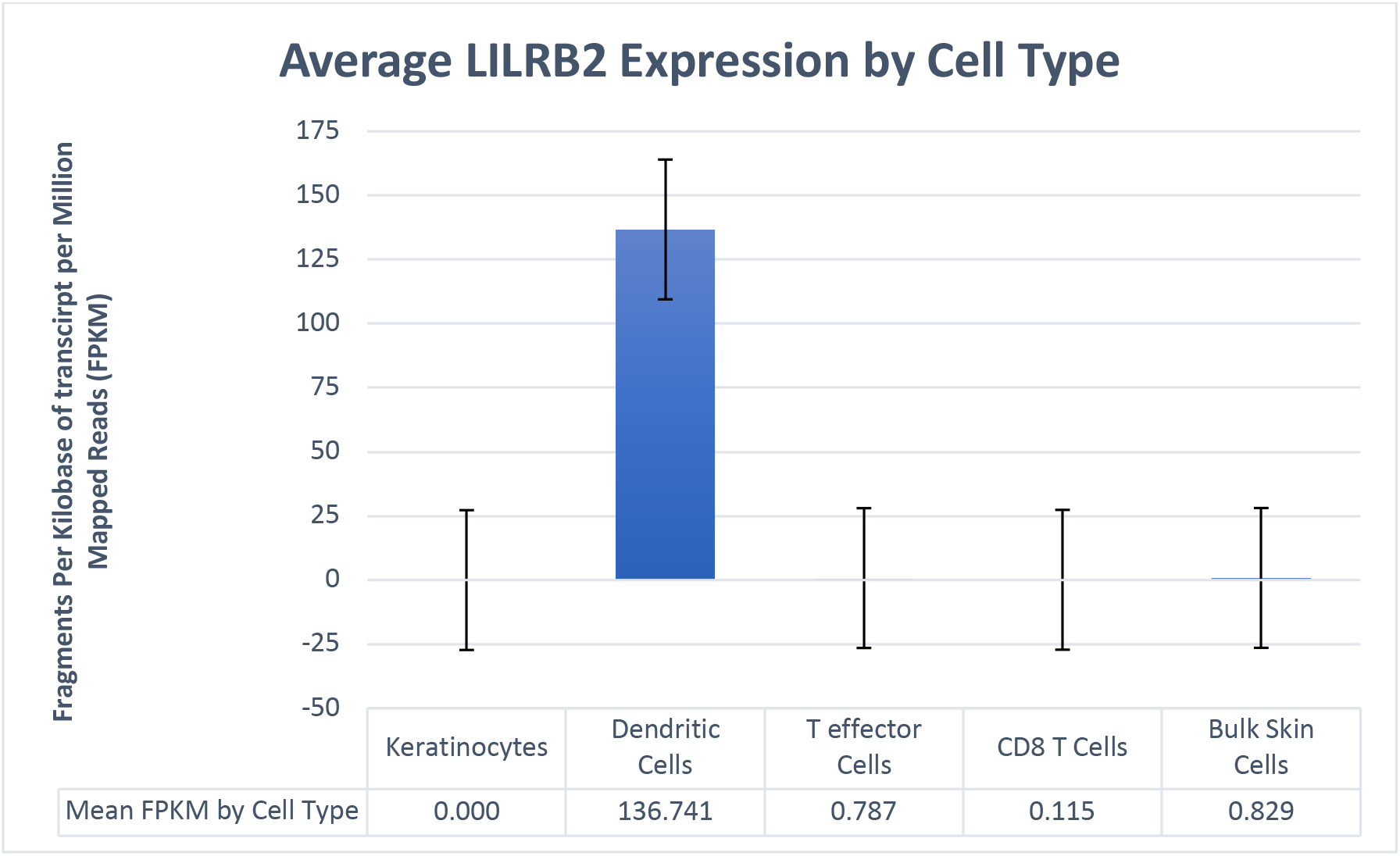
*LILRB2* mRNA is significantly expressed in cutaneous dendritic cells compared to negligible expression in keratinocytes, T cells, and bulk skin cells.

## DISCUSSION

Our results indicate that psoriasis patients, on average, harbor HLA-B alleles with decreased binding affinity to LILRB2. As LILRB2 is an inhibitory receptor, this decreased affinity may lead to a reduction in APC inhibition that results in more potent T cell reactivity. HLA-B*57:01 and HLA-B*27:05 have been associated with psoriasis in multiple studies (Chen et al., 2012, Okada et al., 2014) and these two alleles are among those with the lowest LILRB2 binding affinities (Table S1). The positive signal observed in our study for the binding of LILRB2 to HLA-B but not HLA-C or HLA-A may be explained by the fact that HLA-C is expressed on the cell surface at roughly one-tenth the level of HLA-A and HLA-B (Apps et al., 2015), and by the observation that there is a much smaller range of variation for LILRB2 binding to HLA-A compared to HLA-B.

The role of LILR molecules in the prevention and pathogenesis of diseases is beginning to become more highly recognized. LILRB2 binding to HLA-B has been shown to significantly impact HIV-1 viral control, whereby weaker binding between LILRB2 and HLA-B was associated with better HIV control via increased dendritic cell function (Bashirova et al., 2014). For autoimmune disease, a large genetic study of Takayasu’s arteritis identified genetic epistasis between LILRA3 and HLA-B*52, demonstrating a multiplicative interactive effect on disease susceptibility (Terao et al., 2018). In addition, a genetic study of ankylosing spondylitis found a protective effect of LILRB1, implying that inhibition of dendritic cells through inhibitory LILR molecules may subvert development of autoimmune or inflammatory disease (Majorczyk et al., 2019). Similarly, a study of systemic lupus erythematous patients showed decreased inhibitory activity by LILRB1 on T cells and reduced expression on B cells compared to healthy controls (Monsivais-Urenda et al., 2007). In contrast, the activating LILRA3 receptor has been shown to be increased in both rheumatoid arthritis and multiple sclerosis patients, as well as associated with disease severity in the latter (An et al., 2010, An et al., 2016).

Intriguingly, there appears to be an evolutionary balance between HLA alleles that promote robust immunity against certain infections, but which may also increase the risk of immune-mediated diseases. For example, HLA Class I alleles that have been shown to be protective against HIV-1 are enriched in psoriasis patients (Chen et al., 2012). Similarly, a compound genotype associated with delayed progression of HIV to AIDS (KIR3DS1 plus HLA-B Bw4-80I) also increases psoriasis susceptibly (Chen et al., 2012). Interactions of HLA to certain KIR molecules, such as KIR3DL1, have been shown to be beneficial in control of HIV (Martin et al., 2002), but have also been associated with psoriasis susceptibility (Ahn et al., 2016). Moreover, a variant upstream of HLA-C associated with HLA-C expression was shown to have significant protective effects on the control of HIV; at the same time, this variant was associated with increased susceptibility to Crohn’s disease (Apps et al., 2013).

Here we found another example of the dual effect on infectious control and autoimmune susceptibility for HLA-B and LILRB2. These findings all illustrate a key paradigm in evolutionary genetics: the precarious balance between genetic variants that boost the immune system’s efficacy against infectious disease with those that may overshoot and increase an individual’s susceptibility to autoimmune conditions (Kulkarni et al., 2008). Such opposing genetic forces of infectious versus autoimmune disease pathology may contribute to the differential binding of LILRB2 to HLA molecules.

To our knowledge, our work is the first to identify a role for LILRB2 in psoriasis susceptibility. Our finding, supported by two independent datasets, expands our mechanistic understanding of the role of HLA in immune-mediated diseases, highlighting the importance of APC regulation.

## METHODS

In order to code the binding affinity of LILRB1 and LILRB2 to various HLA alleles as a genetic variable in each individual, we utilized the previously published binding scores of LILRB1 and LILRB2 to 31 alleles of HLA-A, 50 alleles of HLA-B, and 16 alleles of HLA-C (Bashirova et al., 2014, Jones et al., 2011), (**Table S1**). These scores were determined by incubating LILRB1-Fc or LILRB2-Fc fusion proteins with LABScreen HLA class I single antigen beads and measuring the median fluorescence intensity of each LILRB-HLA pairing (Jones et al., 2011). As previously described (Chen et al., 2012), we used HLA*IMP to impute to four-digit resolution HLA class I alleles (-A, -B, and -C) and HLA class II alleles (-DQA1, -DQB1, and -DRB1) of two GWAS cohorts: the WTCCC cohort of 2,178 psoriasis cases and 5,175 controls (Strange et al., 2010) and the GAIN cohort of 1,368 psoriasis cases and 1,348 controls (Nair et al., 2009). We previously confirmed the accuracy of imputation by comparing results to directly genotyped patients and found a concordance rate of 97.4% (Chen et al., 2012). Each individual was assigned a LILRB1 or LILRB2 binding score for HLA-A, -B, and -C that was the sum of the binding scores of that individual’s two HLA-A, -B, or -C alleles to the respective LILRB locus (Bashirova et al., 2014, Chen et al., 2012).

## Abbreviations

MHC: major histocompatibility complex
NK: natural killer cells
APC: antigen presenting cells
LILR: leukocyte immunoglobulin-like receptors
KIR: killer cell immunoglobulin-like receptors
GWAS: genome-wide association study

## Conflicts of Interest

The authors state no conflict of interest.

## Acknowledgements

The GAIN data set was obtained from the database of Genotypes and Phenotypes (dbGaP) found at http://www.ncbi.nlm.nih.gov/gap through dbGaP accession number phs000019.v1.p1. This study makes use of data generated by the Wellcome Trust Case–Control Consortium. A full list of the investigators who contributed to the generation of the data is available at www.wtccc.org.uk. This work was supported by funding from the NIH to W.L. (5U01AI119125).

This project has been funded in whole or in part with federal funds from the Frederick National Laboratory for Cancer Research, under Contract No. HHSN261200800001E. The content of this publication does not necessarily reflect the views or policies of the Department of Health and Human Services, nor does mention of trade names, commercial products, or organizations imply endorsement by the U.S. Government. This Research was supported in part by the Intramural Research Program of the NIH, Frederick National Lab, Center for Cancer Research.

The authors would like to acknowledge Rashmi Gupta for her contributions.

## DATA AVAILABILITY

The GAIN dataset used for the analyses described in this manuscript was obtained from the database of Genotypes and Phenotypes (dbGaP) found at http://www.ncbi.nlm.nih.gov/gap through dbGaP accession number phs000019.v1.p1. Samples and associated phenotype data for the Collaborative Association Study of Psoriasis were provided by Drs. James T. Elder (University of Michigan, Ann Arbor, MI), Gerald G. Krueger (University of Utah, Salt Lake City, UT), Anne Bowcock (Washington University, St. Louis, MO), and Gonc, alo R. Abecasis (University of Michigan, Ann Arbor, MI). This study makes use of data generated by the Wellcome Trust Case-Control Consortium. The Wellcome Trust Case-Control Consortium data were obtained from the WTCCC official website (http://www.wtccc.org.uk/). A full list of the investigators who contributed to the generation of the data is available from www.wtccc.org.uk.

## SUPPLEMENTARY MATERIALS

**Table S1.** HLA Class I allele-specific binding scores of LILRB1 and LILRB2 for viral load controlled determined via univariate model, from (Bashirova et al., 2014)

| HLA Class I Allele | Binding Score<br>LILRB1 | Binding Score<br>LILRB2 |
| --- | --- | --- |
| A*01:01 | 0.36 | 0.55 |
| A*02:01 | 0.11 | 0.60 |
| A*02:03 | 0.17 | 0.38 |
| A*02:06 | 0.28 | 0.52 |
| A*03:01 | 0.41 | 0.51 |
| A*11:01 | 0.48 | 0.48 |
| A*11:02 | 0.63 | 0.80 |
| A*23:01 | 0.38 | 0.51 |
| A*24:02 | 0.43 | 0.55 |
| A*24:03 | 0.41 | 0.98 |
| A*25:01 | 0.11 | 0.56 |
| A*26:01 | 0.11 | 0.67 |
| A*29:01 | 0.18 | 0.56 |
| A*29:02 | 0.13 | 0.53 |
| A*30:01 | 0.56 | 0.75 |
| A*30:02 | 0.21 | 0.40 |
| A*31:01 | 0.24 | 0.48 |
| A*32:01 | 0.10 | 0.58 |
| A*33:01 | 0.07 | 0.58 |
| A*33:03 | 0.12 | 0.66 |
| A*34:01 | 0.14 | 0.57 |
| A*34:02 | 0.26 | 0.50 |
| A*36:01 | 0.20 | 0.40 |
| A*43:01 | 0.07 | 0.39 |
| A*66:01 | 0.26 | 0.75 |
| A*66:02 | 0.20 | 0.53 |
| A*68:01 | 0.13 | 0.43 |
| A*68:02 | 0.11 | 0.43 |
| A*69:01 | 0.18 | 0.65 |
| A*74:01/2 | 0.04 | 0.53 |
| A*80:01 | 0.12 | 0.56 |
| B*07:02 | 0.22 | 0.56 |
| B*08:01 | 0.38 | 0.57 |
| B*13:01 | 0.12 | 0.50 |
| B*13:02 | 0.18 | 0.38 |
| B*14:01 | 0.27 | 0.40 |
| B*14:02 | 0.21 | 0.21 |
| B*15:01 | 0.31 | 0.44 |
| B*15:02 | 0.37 | 0.29 |
| B*15:03 | 0.35 | 0.49 |
| B*15:10 | 0.22 | 0.20 |
| B*15:11 | 0.27 | 0.47 |
| B*15:12 | 0.39 | 0.38 |
| B*15:13 | 0.19 | 0.37 |
| B*15:16 | 0.27 | 0.31 |
| B*18:01 | 0.40 | 0.66 |
| B*27:05 | 0.27 | 0.14 |
| B*27:08 | 0.49 | 0.30 |
| B*35:01 | 0.36 | 0.41 |
| B*38:01 | 0.30 | 0.39 |
| B*39:01 | 0.35 | 0.32 |
| B*40:01 | 0.18 | 0.40 |
| B*40:02 | 0.23 | 0.33 |
| B*40:06 | 0.24 | 0.51 |
| B*41:01 | 0.39 | 0.39 |
| B*42:01 | 0.42 | 0.38 |
| B*44:02 | 0.19 | 0.36 |
| B*44:03 | 0.19 | 0.16 |
| B*45:01 | 0.45 | 0.54 |
| B*46:01 | 0.31 | 0.57 |
| B*48:01 | 0.14 | 0.52 |
| B*49:01 | 0.06 | 0.24 |
| B*50:01 | 0.26 | 0.32 |
| B*51:01 | 0.18 | 0.46 |
| B*51:02 | 0.23 | 0.33 |
| B*52:01 | 0.10 | 0.54 |
| B*53:01 | 0.36 | 0.47 |
| B*54:01 | 0.25 | 0.26 |
| B*55:01 | 0.21 | 0.52 |
| B*56:01 | 0.25 | 0.27 |
| B*57:01 | 0.07 | 0.09 |
| B*57:03 | 0.26 | 0.25 |
| B*58:01 | 0.03 | 0.12 |
| B*59:01 | 0.18 | 0.37 |
| B*67:01 | 0.47 | 0.41 |
| B*73:01 | 0.33 | 0.53 |
| B*78:01 | 0.34 | 0.72 |
| B*81:01 | 0.32 | 0.50 |
| B*82:01 | 0.36 | 0.41 |
| C*01:02 | 0.04 | 0.39 |
| C*02:02 | 0.19 | 0.54 |
| C*03:02 | 0.13 | 0.29 |
| C*03:03 | 0.20 | 0.30 |
| **C\*03:04** | **0.39** | **0.37** |
| **C\*04:01** | **0.04** | **0.18** |
| **C\*05:01** | **0.24** | **0.49** |
| **C\*06:02** | **0.31** | **0.76** |
| **C\*07:02** | **0.15** | **0.62** |
| **C\*08:01** | **0.22** | **0.36** |
| **C\*12:03** | **0.21** | **0.54** |
| **C\*14:02** | **0.16** | **0.5** |
| **C\*15:02** | **0.36** | **0.45** |
| **C\*16:01** | **0.14** | **0.50** |
| **C\*17:01** | **0.08** | **0.27** |
| **C\*18:02** | **0.29** | **0.61** |

